# Spontaneous aggressive ERα+ mammary tumor model is driven by Kras activation

**DOI:** 10.1101/442624

**Authors:** Katie M. Campbell, Kathleen A. O’Leary, Debra E. Rugowski, William A. Mulligan, Erica K. Barnell, Zachary L. Skidmore, Kilannin Krysiak, Malachi Griffith, Linda A. Schuler, Obi L. Griffith

**Author notes:** Senior Authors (Linda A. Schuler, Obi L. Griffith). **Correspondence** (O.L.G.), (L.A.S.).

## Abstract

The NRL-PRL murine model, defined by mammary-selective transgenic rat prolactin ligand *rPrl* expression, establishes spontaneous ER+ mammary tumors, mimicking the association between elevated prolactin (PRL) and risk for development of ER+ breast cancer in postmenopausal women. Whole genome and exome sequencing in a discovery cohort (n=5) of end stage tumors revealed canonical activating mutations and copy number amplifications of *Kras*. The frequent mutations in this pathway were validated in an extension cohort, identifying activating *Ras* alterations in 79% (23/29) of tumors. Transcriptome analyses over the course of oncogenesis revealed marked alterations associated with Ras activity in established tumors, compared to preneoplastic tissues, in cell-intrinsic processes associated with mitosis, cell adhesion and invasion, as well as in the tumor microenvironment, including immune activity. These genomic analyses suggest that PRL induces a selective bottleneck for spontaneous Ras-driven tumors which may model a subset of aggressive clinical ER+ breast cancers.

## Introduction

Epidemiologic evidence has linked higher levels of circulating prolactin (PRL) with increased risk for estrogen receptor alpha (ER+) breast cancers, particularly in postmenopausal women (Tikk et al., 2014; Tworoger and Hankinson, 2008; Tworoger et al., 2013). However, the role of PRL in tumor progression is more controversial (Hachim et al., 2016; Shemanko, 2016; Tworoger and Hankinson, 2008). Moreover, activation of STAT5A, the downstream mediator of canonical PRL signals, is associated with a better prognosis (Peck et al., 2012; Tworoger and Hankinson, 2008). In order to distinguish the contributions of PRL to breast cancer from its actions in pregnancy, we generated the NRL-PRL murine model, where transgenic rat PRL (rPRL) is secreted by mammary epithelia to mimic the local production of PRL reported in women (Marano and Ben-Jonathan, 2014; McHale et al., 2008; O’Leary et al., 2015; Rose-Hellekant et al., 2003). In young adult nulliparous NRL-PRL females, prior to evidence of detectable mammary lesions, ductal epithelia proliferate more rapidly and exhibit increased progenitor activity and Wnt signaling, resulting in reduced maturation of luminal progenitors compared to wildtype (WT) littermates (O’Leary et al., 2017; Rose-Hellekant et al., 2003). With age, NRL-PRL females develop hyperplastic lesions, which strongly express ER and progesterone receptor (PR), and eventually develop histologically diverse and metastatic carcinomas (O’Leary et al., 2015; Rose-Hellekant et al., 2003). In contrast to the preneoplastic lesions, established tumors express variable ER and low PR, resembling the luminal B subtype of clinical breast cancer (Arendt et al., 2011; O’Leary et al., 2015).

In order to understand the mechanisms that underlie the ability of PRL to drive the development of cancers in this model, we used comprehensive genomic analyses to identify genomic alterations and patterns of gene expression associated with cancer progression. In a Discovery Set of five independent, histologically diverse carcinomas and matched adjacent tumor-free mammary glands (AMG) from aged NRL-PRL females, we found that all tumors contained somatic alterations in *Kras*, including canonical activating mutations (i.e., G12, G13, Q61) or amplifications that resulted in elevated Ras pathway activation. *Kras* was not altered in any preneoplastic mammary glands. Our findings were validated in an Extension Set of 22 tumors and 4 cell lines (derived from two additional tumors), demonstrating activating *Ras* alterations in 79% of tumors. Ras activation was associated with increased phosphorylation of ERK1/2 and transcripts for Ras target genes and some pathway mediators, but variable *rPrl* transgene expression and reduced canonical downstream mediators of PRL signaling. These findings coincide with recent reports that elevated Ras signaling drives many clinical luminal B cancers, and is associated with a poor prognosis (Olsen et al., 2017; Wright et al., 2015). In contrast to *Kras*-activated tumors, preneoplastic mammary glands maintained constitutive expression of *rPrl* and displayed elevated transcripts for cytokines associated with pro-tumor myeloid populations. Together, these data indicate that constitutive PRL signaling in mammary glands induces a pro-tumor microenvironment, facilitating Ras pathway activation and spontaneous tumorigenesis.

## Results

To understand the molecular events underlying the development of the prolactin-induced carcinomas in NRL-PRL females, we interrogated a Discovery Set from 10 NRL-PRL female mice, consisting of five young adults and five aged tumor-bearing mice. Mammary cell preparations (MCP) were harvested from caudal glands of NRL-PRL females (n=5) at 12 weeks (young adults), prior to morphologic evidence of pathology. Following the formation of spontaneous tumors at 15-20 months of age, three tissues from these mice were collected: 1) end stage mammary tumors, 2) adjacent non-tumor mammary tissue (aged mammary glands, AMG), and 3) matched tail tissue (**Figure 1A, Table S1**). Mammary glands from 12 week-old mice contained primarily simple ductal structures and expressed ER and PR at levels similar to wildtype (WT) age-matched females (**Figure 2A**). In contrast, mammary glands of aged NRL-PRL females contained hyperplastic epithelial structures and expressed ER/PR similar to ductal structures of young glands of either genotype (**Figure 2B**). The ER+ tumors that develop in this model are histotypically diverse, including more differentiated adenocarcinomas (such as T2, T3) and less differentiated spindle cell carcinomas (such as T1, T4, T5). These end stage tumors expressed lower levels of PR than structures in nondysplastic or hyperplastic AMGs (**Figure 2B**).

**Figure 1.**
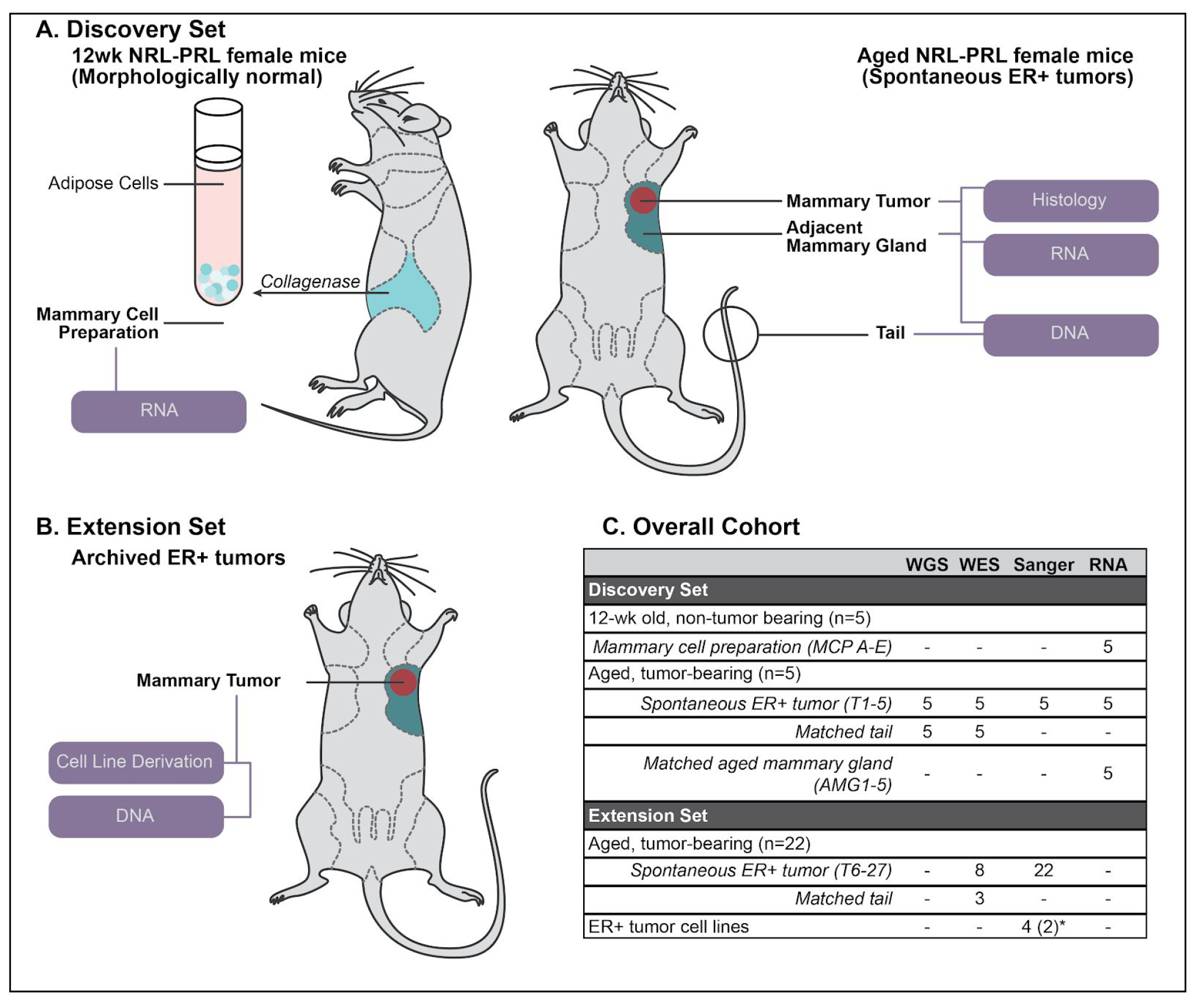
Sample cohort summary. **A.** For our Discovery Set, mammary cell preparations (MCP) were isolated from caudal glands of 12 week old NRL-PRL females (MCP A-E, n=5). Mice with matched end stage tumor (T) and adjacent aged mammary glands (AMG) were collected following the development of spontaneous ER+ tumors (15-20 months; T1-5, AMG1-5, n=5). All samples were examined by RNAseq. Tumors and matched tail DNA samples underwent WGS and WES. **B.** In our Extension Set, we evaluated 22 additional tumors from NRL-PRL females (T6-27) and 4 cell lines (derived from 2 independent tumors). **C.** Summary of samples interrogated in this study. The number of mice associated with each group or the sample identifiers are indicated in parentheses. *Four cell lines were derived from 2 additional independent tumors (2 each).

**Figure 2.**
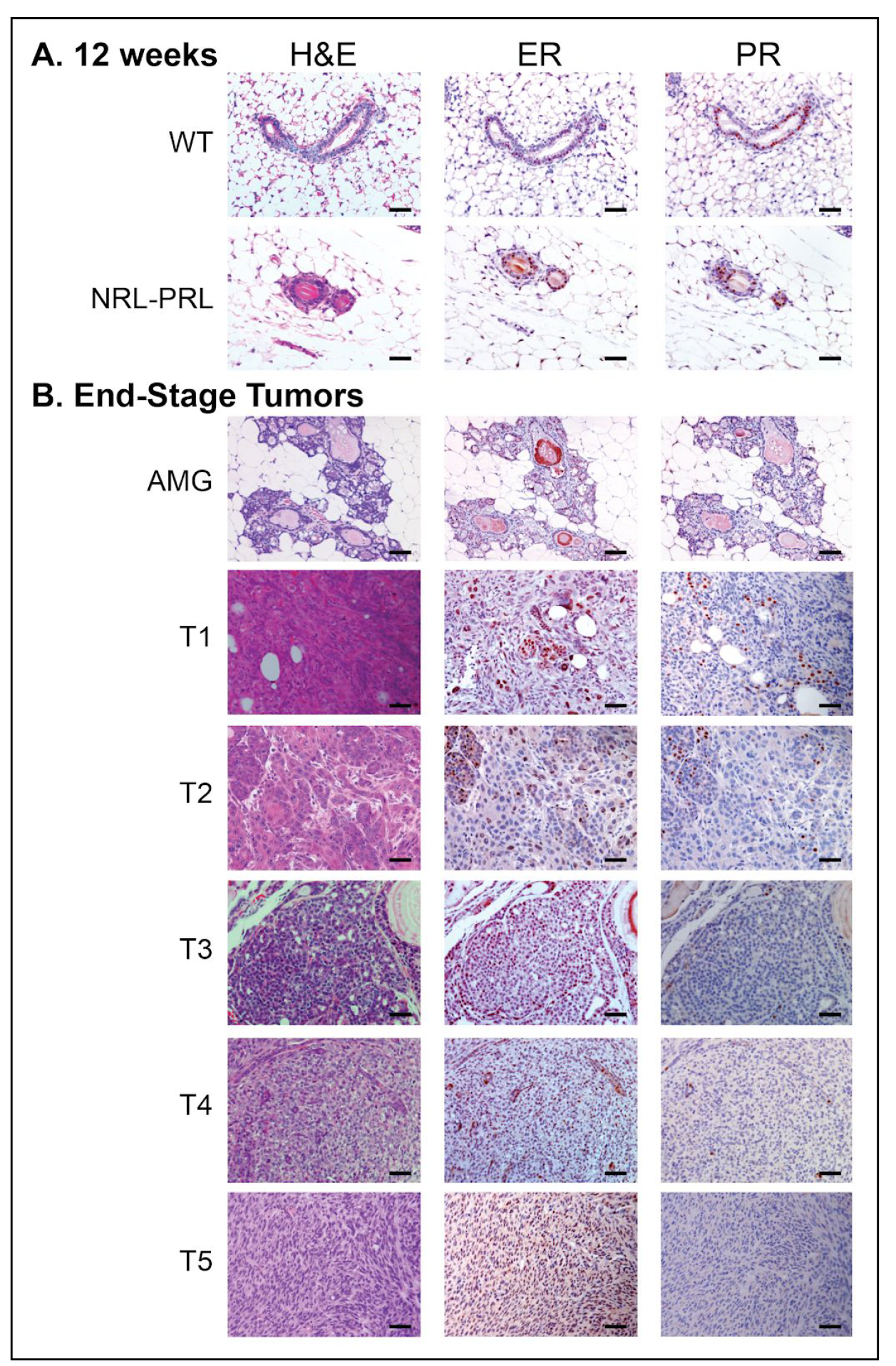
Histology of mammary glands and histologically diverse carcinomas sequenced. **A.** Mammary glands of 12 week old NRL-PRL females did not exhibit marked morphological differences or express different levels of ER and PR from wildtype (WT) age-matched females. **B.** Histology of tumors sequenced and characterized in the Discovery Set and representative adjacent glands. Hematoxylin and eosin (H&E), estrogen receptor alpha (ER), progesterone receptor (PR). Scale bars, 50 microns.

### Genomic characterization

Whole genome and exome sequencing (WGS, WES) was performed on end stage tumors and matched tails for mice within the Discovery Set (n=5). Somatic variants were detected by comparing tumors to matched tail sequencing data. A range of 3-20 nonsilent mutations (median 12) were identified in each tumor (**Figures 3A and S1, Table S2**). *Kras* was the only recurrently mutated gene in this cohort, with missense mutations detected in 4 out of 5 tumors. Activating mutations occurred at two known hotspots (three G12D and one Q61L) and were present among the highest fractions (13-67% DNA VAF) within tumors, suggesting their presence in the founding clone of the tumors. *Kras* activating mutations were confirmed in the RNA of end stage tumors (at 46-75% VAF); however, they were not present in RNA derived from adjacent AMG tissues. The tumor without an activating mutation (T5) was assessed for copy number and structural variants of *Kras*, and a focal amplification and structural variant were identified containing the *Kras* gene locus. There were no large scale copy number alterations, structural variants, or gene-gene fusions recurrently altered across the Discovery Set outside of *Kras* (**Figure S1**).

**Figure 3.**
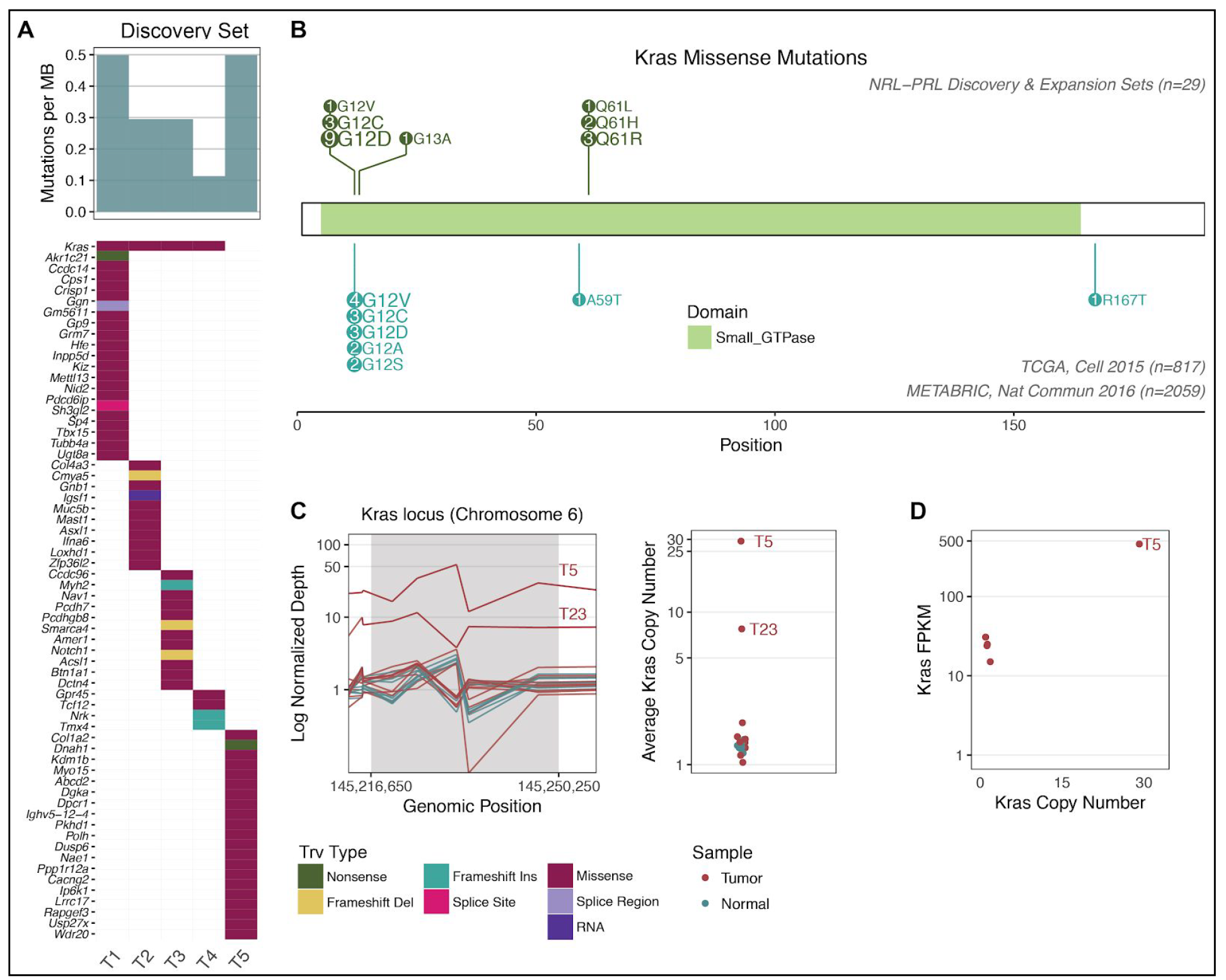
Summary of Kras modifications in NRL-PRL tumors. **A**. Mutation burden ranged from 8-28 nonsynonymous coding mutations across the five tumors in the Discovery Set (median 18; top barchart). The only recurrently mutated gene in our cohort was *Kras* (top row of bottom plot). The waterfall plot indicates the types of mutations detected in the remainder of the cohort (as detailed in the legend below panel C). **B.** Lolliplot of all mutations, comparing the prevalence of *Kras* missense mutations in the tumors in NRL-PRL mice (n = 29; 20 contained *Kras* missense mutations) to *KRAS* missense mutations in human breast cancer (TCGA, n=817; METABRIC, n=2,059). **C.** In the left panel, points represent the normalized depth of sequencing (coverage normalized by respective sample total sequencing depth) over the exons spanning the *Kras* gene locus (in 320-920 bp windows) of all tumor and normal samples. Tumor samples are not normalized with respect to matched normal data, since matched normal tails were not available for all mice. In the right panel, the average depth of the points within the *Kras* locus is indicated on a log2 scale. Amplifications detected in T5 and T23 are labeled. **D.** RNAseq data were available for T1-5 (Discovery Set). The copy number (right panel of C) of *Kras* is plotted along the x-axis, and the gene expression of *Kras* (in fragments per kilobase transcript per million mapped reads, FPKM; quantified by Cufflinks, see Methods) is indicated on the y-axis.

### Confirmation of Kras activation in *rPrl*-induced tumors

We hypothesized that prolonged PRL exposure was creating a selective bottleneck for activating *Kras* mutations in the NRL-PRL mouse model. To determine whether mutations at this locus were consistent in a larger set of these tumors, Sanger sequencing was performed on *Kras* exons 2 and 3 (containing the G12, G13, and Q61 hotspot loci) in an Extension Set of 22 independent PRL-induced tumors (15 archival tumors, T6-T20, and 7 fresh tumors with matched tails, T21-T27) and 4 cell lines derived from 2 additional PRL-induced tumors (**Figure 1B, Table S1**). Hotspot mutations were identified in 14 tumors and all cell lines in this set (**Figure 3B, Table 1**). Overall, the most common alteration was *Kras* G12D (n=9). This is consistent with the mutational activation of *KRAS* in human breast cancers in the TCGA and METABRIC breast cancer datasets, where amino acid G12 is the most recurrently mutated position (**Figure 3B**). Amino acid Q61 was not mutated in any cases from either of these human breast cancer patient cohorts (Cancer Genome Atlas Network, 2012; Curtis et al., 2012).

**Table 1.**
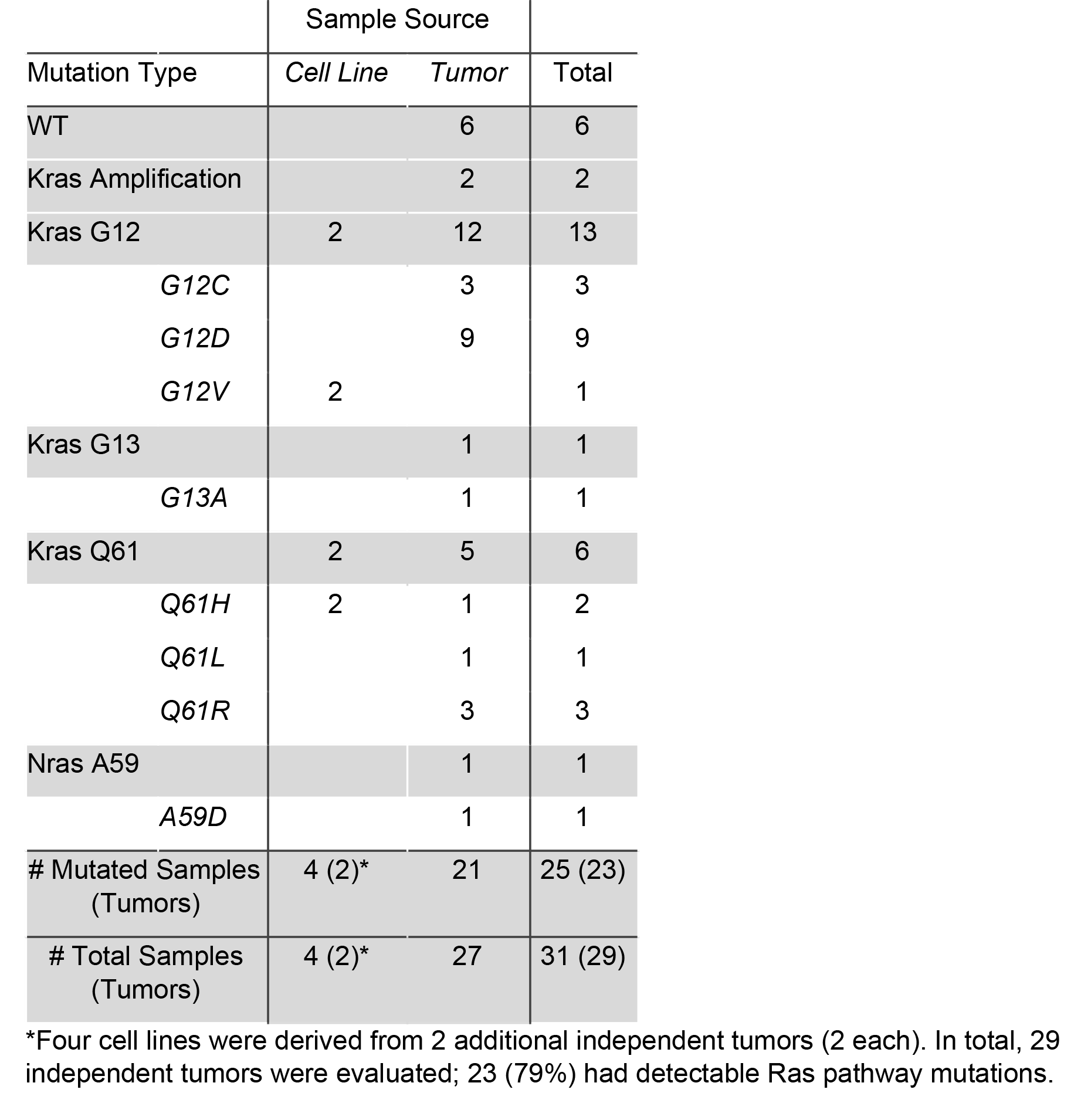
Summary of *Ras* Alterations in the Discovery and Extension Sets.

WES was performed on the eight tumors within the Extension Set that did not contain *Kras* activating mutations in exons 2 or 3. The mutational burden in these tumors ranged from 0-12 (median 6). Other mutations were not identified in *Kras*; however, there was a missense mutation in *Nras* in one tumor (T21, A59D; 21.3% VAF, **Table S2**). The copy number landscape was evaluated for *Kras* copy number alterations, as identified in T5 from the Discovery Set. Copy number was evaluated by quantifying the depth of exome sequencing and normalized per sample. A putative copy number increase in *Kras* was identified in one other tumor (T23), with an average copy number of 7.76 (**Figure 3C**). For reference, the average copy number within the Discovery Set was 1.34, whereas T5 exhibited an average copy number of 29. Given that increased copy number is significantly associated with higher *Kras* expression in T5, we hypothesized that copy number alterations in sample T23 could impact transcript expression (**Figure 3D**). Together, these data demonstrated that at least 23 of 29 (79%) independent spontaneous NRL-PRL tumors developed alterations in Ras genes. We were unable to identify similar somatic alterations using WES in 6 tumors from the Extension Set (T6, T7, T12, T13, T17, T25). Five of these tumors did not have matched normal tails available, reducing confidence in somatic mutation calling and filtering. Additional WGS of these tumors may provide further resolution of the somatic landscape. We focused our subsequent expression analysis on the Discovery Set, which like the majority of the tumors, had *Kras* activating alterations.

### Ras pathway activation differentiates expression in tumors and dysplastic glands

Gene expression was evaluated across the samples from the Discovery Set, including five nondysplastic MCPs from young females, and dysplastic AMGs and end stage tumors from five aged tumor-bearing mice. Unsupervised approaches indicated clustering based upon sample source, where end stage tumors behaved more similarly to other tumors than their matched, adjacent AMGs (**Figure 4A**). In order to understand the varied expression patterns across sample types, we first quantified *rPrl* transgene expression in each sample. *rPrl* expression was maintained in both nondysplastic MCPs and dysplastic AMGs, but varied in tumors (**Figure 4B**). Lower grade, more differentiated tumors (T2 and T3) contained relatively high levels of transgene transcripts, whereas high grade spindle cell carcinomas (T1, T4 and T5) expressed very low levels of *rPrl* mRNA. This association suggests that *rPrl* transgene expression may influence tumor phenotype, but is not required for tumor maintenance.

**Figure 4.**
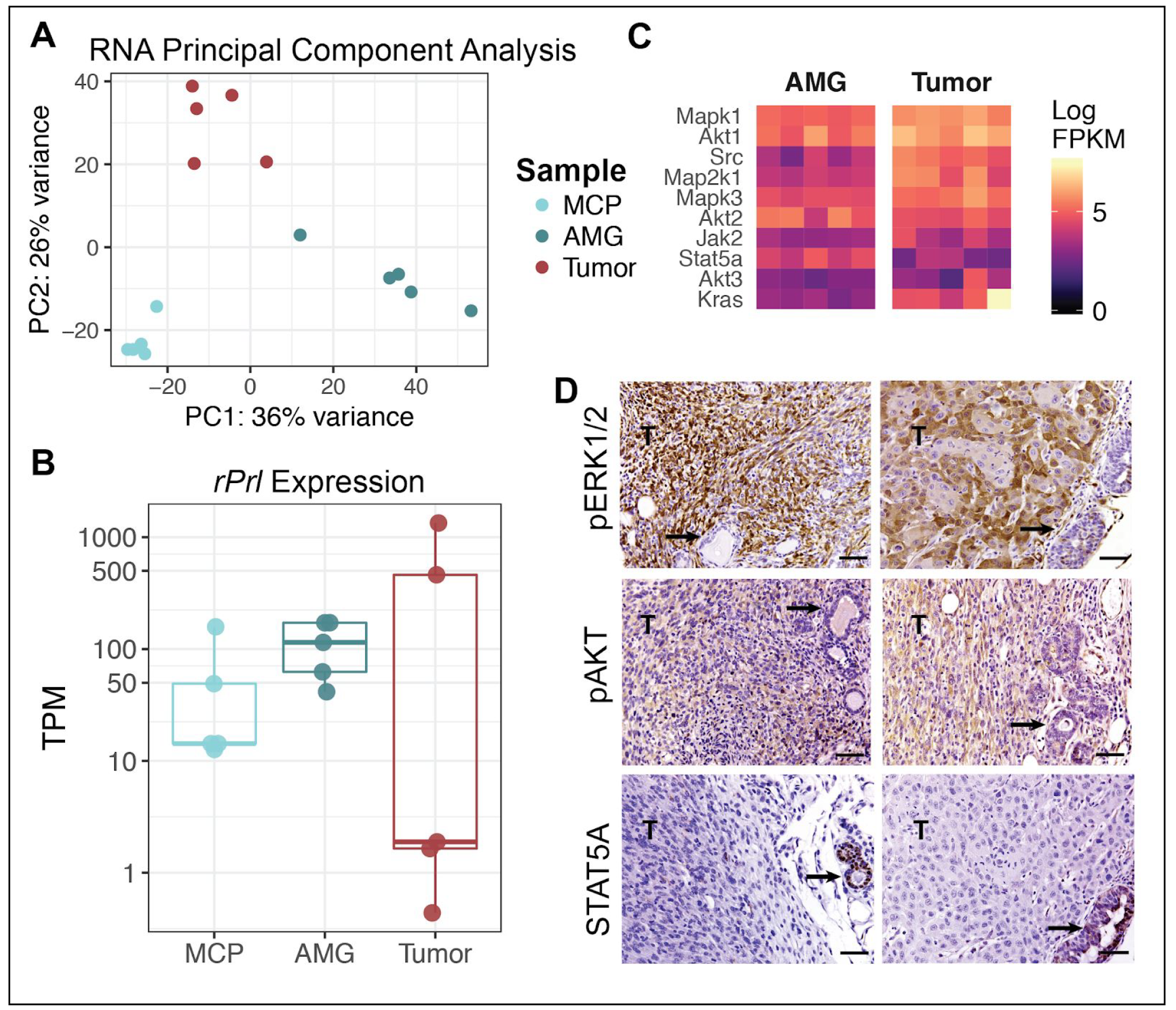
Constitutive prolactin signaling induces a bottleneck for Ras pathway activation. **A.** Unsupervised principal component analysis was performed across nondysplastic MCPs (n=5), aged glands (AMG; n=5), and end stage spontaneous ER+ tumors (n=5) from the Discovery Set. Color indicates sample type. **B.** Kallisto was used to quantify transgene expression in transcripts per million (TPM). Pseudoalignment of RNA reads was performed against the annotated mouse transcriptome and *rPrl* (ENSRNOT00000023412.4). **C.** Heatmap displaying expression of canonical downstream mediators of prolactin signaling (*Jak2, Stat5a*) and Ras pathway mediators (*Mapk1/3, Akt1/2/3, Src, Map2k1, Kras*). Fill indicates expression level in FPKM for AMG and tumor samples from the Discovery Set. **D.** Immunohistochemical staining for STAT5A, pERK1/2, and pAKT in two representative tumors of different histotypes with adjacent glands (left, right). Tumor regions are indicated by ‘T’ in the image, and arrows point to adjacent AMGs. Scale bars, 50 microns.

To determine the downstream effects specific to *Kras* activation, supervised differential expression analysis was performed, comparing matched tumors to adjacent AMGs. There were 41 genes that were differentially expressed between these two tissue types (q<0.001) that were also annotated as part of the KEGG Ras pathway (mmu04014). This included upregulation of *Mapk1, Map2k1*, and *Src* mRNAs in end stage tumors (**Figure 4C**), indicating higher levels of transcripts for mediators of the Ras pathway in *Kras*-driven tumors than AMGs. Consistently, tumors, but not adjacent AMGs, displayed strong phosphorylation of ERK1/2 and AKT by immunohistochemistry (**Figure 4D**).

In normal mammary function, most PRL signals are mediated by the JAK2/STAT5A pathway (Oakes et al., 2008). Activation of this signaling cascade results in the phosphorylation and dimerization of STAT5A, which then translocates to the nucleus to regulate transcription (Hammarén et al., 2018). Interestingly, *Stat5a* mRNA was significantly reduced in end stage tumors, compared to AMGs (q=1.54e-5, **Figure 4C**), particularly in poorly differentiated tumors T1, T4, and T5. Nontumor mammary structures adjacent to the tumors also displayed strong nuclear STAT5A staining (**Figure 4D**), suggesting that the constitutive *rPrl* expression seen consistently across AMGs was activating this signaling cascade.

### Tumorigenic processes and the microenvironment differ across stages of disease progression

Subsequent pathway analysis using the RNAseq data from the Discovery Set revealed multiple changes with respect to disease progression. Increases in tumorigenic processes, including cell cycle regulation and cell adhesion, were associated with Ras pathway activation in end stage tumors (**Figure 5A-B**). Chromosome organization and chromatin structure, mitotic nuclear division, as well as cadherin and proteins involved in cell adhesion and adherens junctions, were significantly upregulated in tumors compared to AMGs or MCPs (**Figure 5B**). This is consistent with increased mitotic and invasive properties of cancer cells. AMGs displayed increased expression of genes related to fatty acid metabolism, relative to MCPs and tumors (p<0.01, **Figure 5A-B**), reflecting the higher proportion of adipocytes in these preparations.

**Figure 5.**
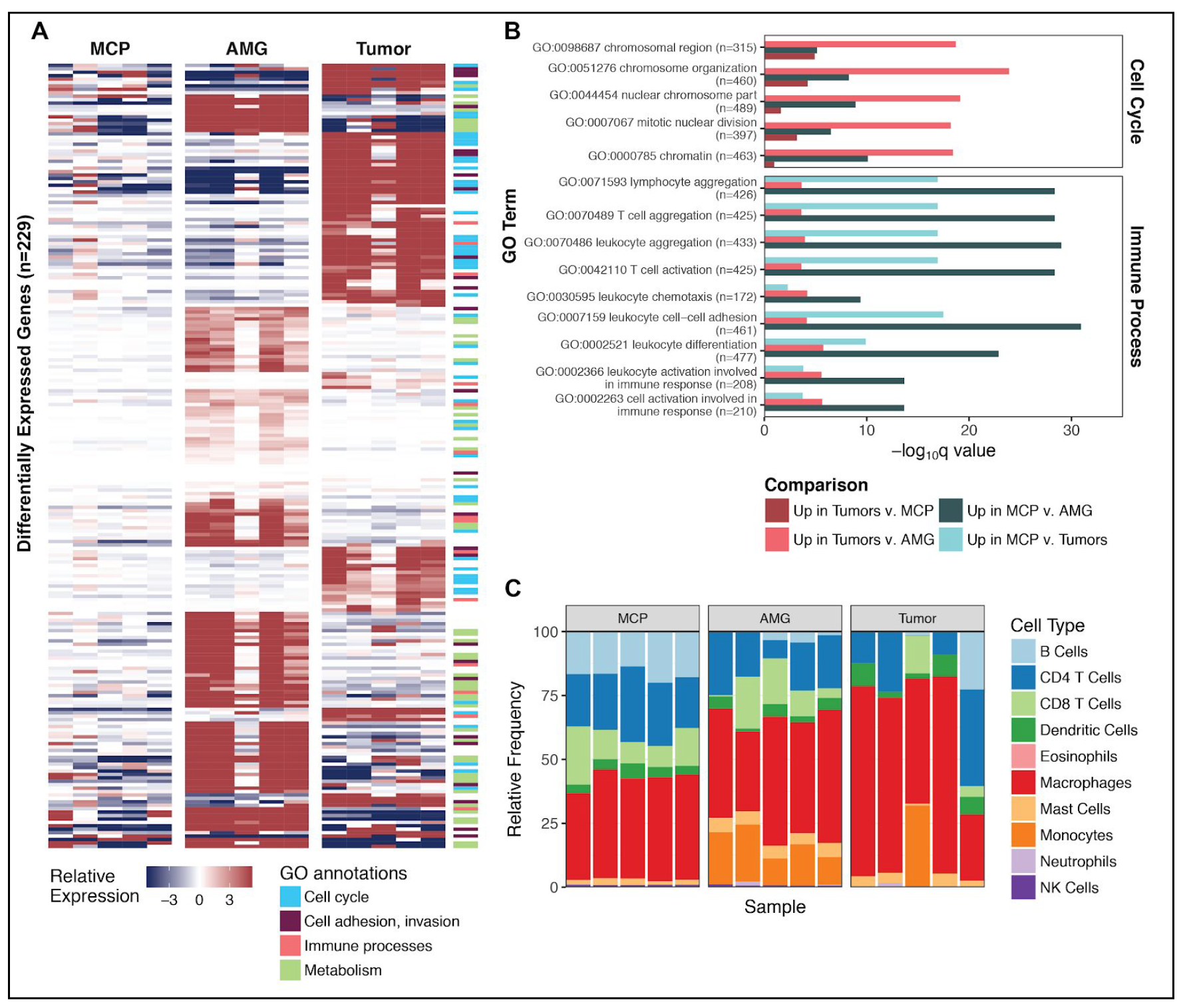
Tumorigenic processes differ across stages of disease progression. **A.** Supervised differential expression analysis was performed on all comparisons of nondysplastic MCPs (n=5), AMGs, and end stage tumors (n=5) from the Discovery Set. Genes that were significantly upregulated (p<0.0001) in either tumors (compared to AMGs and MCPs) or glands (compared to tumors and MCPs) that were also annotated within significantly upregulated GO annotations (p<0.001) are shown in the heatmap (n=229). Fill indicates the gene-median centered expression value, calculated with respect to the entire Discovery Set. GO annotations were grouped into cellular processes, and genes corresponding to each annotation are noted in the side color bar (see Methods). **B.** A representative set of differentially expressed GO process, categorized as ‘Cell Cycle’ and ‘Immune Processes’ in Panel A, are displayed based upon their q value. Following each GO process, the number of genes included in the annotation is indicated. Comparisons are summarized as those that were upregulated in either MCPs or Tumors, compared to the other sample types. **C.** seq-ImmuCC was used to predict the relative proportions of immune cell populations (y axis) in MCPs, AMGs, and end stage tumors from the Discovery Set. Samples are ordered by identifier (MCP A-E, AMG 1-5, and Tumors 1-5).

Compared to MCPs, both AMGs and tumors showed significantly decreased expression in processes related to immune activation, including leukocyte cell-cell adhesion, leukocyte and lymphocyte aggregation, and T cell activation and aggregation (q<1e-29; **Figure 5A-B**). These observations led us to hypothesize that the immune microenvironment differed across stages of disease. Markers of hematopoietic and immune cell lineages were significantly downregulated in tumors compared to preneoplastic cells, including *Ptprc, Cd8a, Cd4, Tra, Trb, Cd40*, and *Cd19* mRNAs (p<0.001), suggesting that tumors have reduced immune infiltrate, specifically those responsible for adaptive immune recognition and rejection (**Table S3**). We therefore interrogated the RNAseq data to predict the relative proportions of infiltrating immune cell subpopulations using deconvolution approaches to further resolve the immune landscape across the Discovery Set (Chen et al., 2018). All samples displayed a mixture of immune cell subpopulations, and the distinct tumors showed considerable diversity in the predicted proportions of these immune cell types (**Figure 5C**). Macrophages represented the predominant immune cell population across all samples (**Figure 5C**); however, these algorithmic approaches only determined the relative proportion of immune cell subpopulations, without quantifying the absolute levels of immune infiltrate in each sample.

In order to confirm findings and evaluate trends in the RNAseq data, we examined select transcripts by RT-PCR in the Discovery Set. In addition, MCPs were generated from WT 12 week old females (WT MCPs) and from aged NRL-PRL females (AMCPs), enabling us to assess processes associated with preneoplastic changes in similar mammary cell preparations. Lipid metabolic enzymes (*Acadm, Fasn* mRNAs) were significantly upregulated in AMGs compared to AMCPs, confirming that the AMG preparations of tissue adjacent to tumors were enriched with adipocytes (**Figure 6A**).

**Figure 6.**
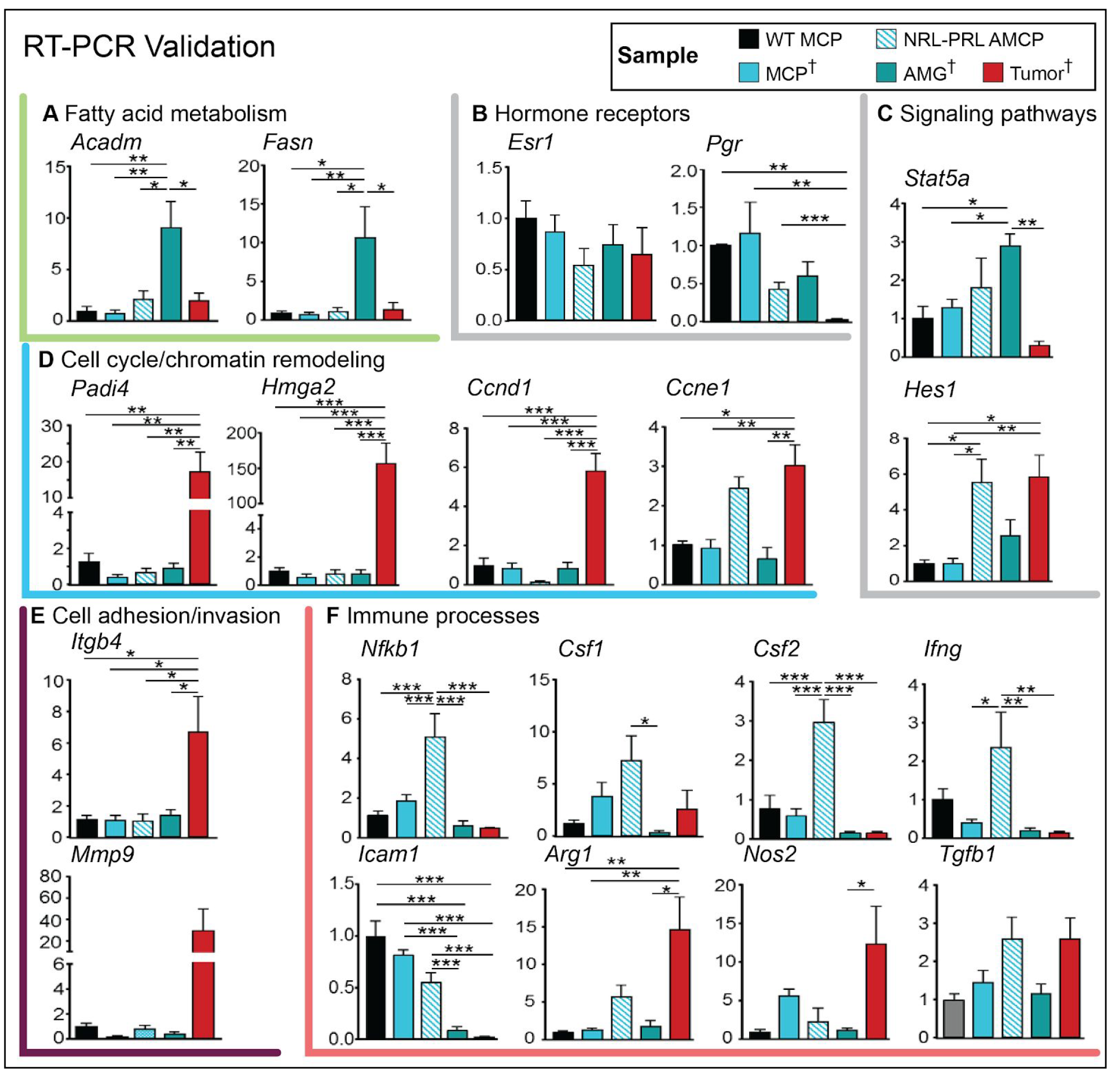
Validation of RNAseq findings. RT-PCR was used to evaluate transcripts associated with cellular processes (A-F) altered with oncogenesis identified in Figure 5 across samples in the Discovery Set (MCP, AMG, Tumor; indicated by †), as well as aged mammary cell preparations from NRL-PRL females (AMCP). MCPs were also isolated from wildtype young adults (WT MCP) (n=4-5). Data are represented as mean +/-S.E.M. Significant differences were determined by one-way ANOVA, followed by the Tukey post-test, ^*^ (p<0.05), ^**^ (p<0.01), and ^***^ (p<0.001).

In order to further elucidate the stages of PRL-induced pathogenesis, hormone receptors and signaling pathways were interrogated. While transcripts for ERα (*Esr1*) were not altered across samples, *Pgr* transcripts fell markedly with disease progression (**Figure 6B**), confirming detected protein expression (**Figure 2)**.

Acquisition of somatic alterations in *Kras* in tumors was associated with loss of *Stat5a* mRNA (**Figure 6C**), consistent with **Figure 4C,D**. Increased Notch activity, indicated by *Hes1* transcripts, and *Ccne1* mRNA in both AMCPs and tumors were consistent with increased proliferation of preneoplastic epithelial populations and tumors in NRL-PRL females (Zender et al., 2013). However, *Ccnd1, Padi4*, and *Hmga2* transcripts were upregulated only in tumors, indicating high proliferation and chromatin remodeling, well-recognized cell autonomous effects of elevated Ras activity (**Figure 6D**). Unsurprisingly, tumors also showed striking elevations in transcripts associated with invasion and interaction with the extracellular matrix (*Itgb4, Mmp9*) (**Figure 6E**).

Interrogation of cytokines and immune mediators revealed significant differences between tumors and preneoplastic mammary tissue (**Figure 6F**). Hyperplastic AMCPs, prior to *Kras* alterations, displayed elevated levels of *Nfkb1, Csf1, Csf2, Ifng*, and *Tgfb1* mRNAs. However, most of these transcripts, along with *Icam1* mRNA, were precipitously lower in established tumors, depicting an immunosuppressed environment. *Tgfb1* remained elevated, and *Arg1* and *Nos2* transcripts were increased in tumors, consistent with shifts in activation of resident myeloid populations, including macrophages and myeloid-derived suppressor cells (Sica and Bronte, 2007). Together, these data support marked changes in the immune microenvironment in PRL-induced mammary pathogenesis.

## Discussion

Increased local exposure to PRL in the NRL-PRL model results in spontaneous diverse ER+ mammary cancers after a long latency, mimicking clinical luminal B, ER+ cancers in postmenopausal women. Here we demonstrated that constitutive PRL signaling creates a pro-tumor niche and an evolutionary bottleneck for Ras pathway activation, particularly *Kras* activating mutations. Elevated Ras activity was associated with reduced activity in the canonical PRL-JAK2/STAT5A signaling cascade despite continued transgene expression in some tumors. Transcriptome analyses over the course of oncogenesis revealed marked alterations associated with Ras activity in established tumors in cell-intrinsic processes associated with mitosis, cell adhesion and invasion, as well as in the tumor microenvironment, including immune activity.

Mirroring the connection to human disease (Tikk et al., 2014; Tworoger and Hankinson, 2008; Tworoger et al., 2013), PRL has been observed to promote mammary carcinogenesis in multiple genetically engineered mouse models (Arendt and Schuler, 2008). Elevated PRL accelerates tumor development in combination with other oncogenes (Arendt et al., 2006, 2009; O’Leary et al., 2014), and conversely, its absence increases tumor latency in combination with viral oncogenes (Oakes et al., 2008; Vomachka et al., 2000). Interestingly, BALB/c mice implanted with pituitary isografts, which elevate circulating PRL, develop mutagen-induced mammary tumors that also exhibit mutations in *Kras (Guzman et al., 1992)*. Furthermore, expression of transgenic *Kras* G12V in beta lactoglobulin-expressing mammary cells during lactation (driven in part by high PRL) results in ER+ tumors in C57BL/6 mice (Andò et al., 2017). The link between PRL and development of Ras-driven mammary tumors in these distinct mouse models in different genetic backgrounds points to strong complementarity of these signals in mammary oncogenesis.

Germline ablation of *Stat1* in 129/SvEv mice results in mammary tumors that acquire a truncating mutation in *Prlr*, resulting in its constitutive activation (Griffith et al., 2016). The well-differentiated tumors in *Stat1*-/-mice are characterized by strong STAT5 phosphorylation (Chan et al., 2014), similar to tumors that develop in response to transgenic *Stat5a* (Iavnilovitch et al., 2004). However, in NRL-PRL females, the robust STAT5 activation in preneoplastic mammary structures was markedly absent in established tumors, associated with reduced *Stat5a* mRNA and acquisition of Ras mutations, demonstrating a transition from the JAK2/STAT5 axis to the Ras signaling pathway. Consistently, tumors that arise in NRL-PRL females lose dependence on JAK2 signaling with disease progression (Sakamoto et al., 2010), whereas mammary tumors in *Stat1*-deficient mice remain dependent on JAK2 (Chan et al., 2014). Together, these data are congruent with a model whereby JAK2/STAT5A signals promote early tumorigenesis, but their ongoing activity supports well-differentiated less aggressive luminal cancers which are more likely to respond to anti-estrogen therapies (Peck et al., 2012; Rädler et al., 2017).

Although all tumors in the NRL-PRL Discovery Set developed alterations in *Ras* family members, any ongoing role for the JAK2-STAT5 pathway and PRL-initiated signals appeared to differ across individual tumors. In the Discovery Set, the more differentiated adenocarcinomas (T2 and T3) maintained transgenic *rPrl* expression, and showed slightly higher *Stat5a* mRNA than the less differentiated spindle cell carcinomas (T1, T4, T5). Continued STAT5A signals may promote cell adhesion and other differentiated characteristics in these tumors (Nouhi et al., 2006; Sultan et al., 2005), despite the overall reduction of *Stat5a* transcripts compared to preneoplastic cells. PRL is able to initiate a diverse spectrum of signals apart from JAK2/STAT5, including activation of Src and downstream pathways, and crosstalk with the Ras cascade itself (Barcus et al., 2013; Erwin et al., 1995; Martín-Pérez et al., 2015). These observations likely account for the controversy surrounding the role of PRL signals in clinical disease (Hachim et al., 2016; Shemanko, 2016).

Analysis of differentially expressed genes and associated processes with disease progression revealed complex alterations in cell signaling, the cytokine milieu, and immune microenvironment early in PRL-driven pathogenesis. Together with PRL-augmented mammary epithelial progenitor cell subpopulations (O’Leary et al., 2017), our findings predict elevated inflammatory cytokines and changes in immune cell subpopulations prior to acquisition of Ras pathway mutations that may contribute to tumorigenesis. End stage tumors with genetic alterations in *Kras* exhibited marked modulation of the immune response, including reduced expression of inflammatory cytokines and increased markers of activated myeloid cells. These findings are consistent with the literature describing immune evasion by Kras activation by multiple mechanisms, including reduced expression of tumor antigens, and promotion of recruitment and differentiation of immune populations that inhibit anti-tumor immune activity (Cullis et al., 2017; Dias Carvalho et al., 2018). Future studies will elucidate interactions between preneoplastic/malignant epithelial cells and the immune microenvironment through stages of PRL-induced pathogenesis.

The low rate of somatic mutations and predicted low tumor infiltrating lymphocytes in NRL-PRL tumors mimic the relative immune silence of clinical luminal breast cancers (Dieci et al., 2016; Luen et al., 2016; Vonderheide et al., 2017). Although the frequency of RAS mutations is relatively low in human breast cancer (<5%) (Cancer Genome Atlas Network, 2012; Curtis et al., 2012), recent studies have revealed that a high proportion of luminal B ER+ breast cancers display elevated Ras pathway activity as a result of reduced expression or somatic loss of RasGAP tumor suppressors (Griffith et al., 2018; Olsen et al., 2017; Wright et al., 2015). RAS itself has been an elusive therapeutic target (McCormick, 2015; Tran et al., 2016). However, targeting RAS-modulated immune regulators (Candido et al., 2018; Mitchem et al., 2013) may intercept T cell-suppressive signaling by pro-tumor myeloid populations. This approach may facilitate responses to T cell-directed therapies, such as the recent exciting anecdotal report of T cell transfer in metastatic breast cancer (Zacharakis et al., 2018).

For many years, it was questioned whether mouse models of mammary cancer could be estrogen responsive and model clinical luminal breast cancers. However, the ER+ cancers that develop in the NRL-PRL model, like the *Stat1-/-* and *Kras* (G12V) models (Andò et al., 2017; Chan et al., 2012), demonstrate that the tools available for murine systems can be valuable to study ER+ breast cancer. Even in PRL-induced tumors which have lost dependence on estrogen for growth, ER-mediated signals continue to modulate cell phenotype and cancer stem cell activity (Shea et al., 2018). These preclinical immunocompetent models of estrogen responsive mammary tumors can provide insight into the biological behavior of these cancers, including metastasis, tumor cell plasticity and cancer stem cell activity, therapeutic responsiveness, and patterns of resistance. Our studies of tumorigenesis in the NRL-PRL model describe the ability of constitutive *rPrl* expression to create a pro-tumor mammary microenvironment, converging into a bottleneck conducive for Ras pathway activation, which initiates aggressive tumor growth, invasion, and immunosuppression of the tumor microenvironment, features shared with a subset of clinical ER+ breast cancers.

## Experimental Procedures

### Mice

NRL-PRL mice [lines 1647-13, TgN(Nrl-Prl)23EPS; 1655-8, TgN(Nrl-Prl)24EPS] were generated and maintained in the FVB/N strain background (O’Leary et al., 2015; Rose-Hellekant et al., 2003). All tumors in the Discovery Set were TgN(Nrl-Prl)23EPS; the Extension Set included archived tumors of both lines. Mice were housed and handled in accordance with the Guide for Care and Use of Laboratory Animals in AAALAC-accredited facilities, and all procedures were approved by the University of Wisconsin-Madison Animal Care and Use Committee.

### Sample Acquisition

We examined changes with age and pathogenesis in heterozygotic nulliparous females. Mammary epithelial cells were isolated from 12 week old animals, prior to evidence of morphological abnormalities. Other animals were aged until the primary mammary tumors were 1.5 cm in diameter (end stage), when the tumors, adjacent mammary glands, and tails were collected and flash frozen for subsequent analyses. For the Extension Set, an additional 22 tumors and 2 pairs of cell lines derived from 2 independent tumors were examined. These additional tumors included 7 freshly isolated tumors and matched tails, and 15 archived tumors equally distributed among the glandular, papillary and spindle cell carcinoma histotypes.

### Morphological and Immunohistochemical Analyses

Samples were collected and fixed overnight in 10% neutral buffered formalin and processed into 5 micron sections. Mammary structures were assessed in hematoxylin and eosin (H&E) stained tissue sections, and protein expression was visualized by immunohistochemistry as previously described (Arendt et al., 2011) using antibodies against the following antigens: ER, #SC-542, Santa Cruz Biotechnology (1:500); PR, A0098, Dako (1:500); pAKT^S473^, #3787, Cell Signaling Technologies (1:50); pERK1/2, #9101, Cell Signaling Technologies (1:400); STAT5A, #SC-1081, Santa Cruz Biotechnology (1:5000).

### Library Construction and Sequencing Strategy

Genomic DNA was isolated using the Qiagen Blood & Cell Culture DNA Mini Kit (#13323; Hilden, Germany). Whole genome sequencing (WGS) libraries for matched tumor and normal tail samples (n=5) were prepared using the Illumina TruSeq PCR free kit with dual indexed adaptors. WGS libraries were pooled and sequenced across 1 flow cell (8 lanes) on the Illumina HiSeq X platform with paired 2 × 150 bp reads with median 33.35X coverage (27.18-37.52X). Whole exome sequencing (WES) libraries for matched tumor and normal tail samples were pooled and sequenced on 1 lane of the Illumina HiSeq 4000 with paired 2 × 150 bp reads with median 37.99X coverage (10.39-85.45X). Total RNA was isolated using the Qiagen RNeasy Midi Kit (#75142; Hilden, Germany). RNA sequencing (RNAseq) libraries were prepared using the Illumina TruSeq stranded total RNA kit. Libraries were pooled and sequenced across 4 lanes of a HiSeq 4000 with 2 × 150 bp reads with median 60,072,344 reads (43,482,392-100,068,565 total reads).

### Sanger Validation Sequencing

For the Extension Set and positive controls from the Discovery Set, the regions surrounding *Kras* hotspots G12/G13 and Q61 were amplified from 5ng of genomic DNA, using Phusion Hot Start Flex DNA polymerase (#M0535; New England Biolabs), and primers which tagged products with M13F(−21) (forward strand) and M13R (reverse strand) sequences (see Table S5 for Primer Sequences). PCR products were purified using the QIAquick PCR Purification Kit (#28104; Qiagen), and sent to GeneWiz for Sanger Sequencing. Traces were manually assessed for variants within G12, G13, and Q61 using the CodonCode Aligner software (version 7.0.1; CodonCode Corporation).

### DNA sequencing and analysis

The Genome Modeling System (GMS) was used for all analysis, including the somatic variant detection and RNA-seq analysis (Griffith et al., 2015). WGS and WES data was processed through SpeedSeq v0.1.0 (Chiang et al., 2015; Faust and Hall, 2014), which aligns reads by BWA-MEM v0.7.10 (Li, 2013) to the mouse reference genome (NCBI build 38, GRCm38) and marks duplicates using SAMBLASTER v0.1.22 (Faust and Hall, 2014). Matched (normal) tail samples were analyzed for all 5 tumors in the Discovery Set and 3 tumors in the Extension Set which did not exhibit *Kras* G12, G13, or Q61 mutations. Somatic events were identified by individually comparing tumor DNA to matched tail (normal) DNA. Single nucleotide variants (SNVs) were detected by the union of SomaticSniper v1.0.4 (Larson et al., 2012), VarScan2 v2.3.6 (Koboldt et al., 2012), Strelka v1.0.11 (Saunders et al., 2012), and Mutect v1.1.4 (Cibulskis et al., 2013). Small insertions and deletions (indels) were detected by taking the union of GATK Somatic Indel Detector (v5336) (McKenna et al., 2010), Varscan2, and Strelka. 5 tumors from the Extension Set which did not exhibit *Kras* G12, G13, or Q61 mutations did not have DNA from matched normal tail samples (T6, 7, 12, 13, and 17). For these tumors, SNV/indels were detected by comparing the aligned tumor WES data to the mouse reference genome GRCm38 using Varscan 2 (v2.3.6) (Koboldt et al., 2012).

SNV/indels were filtered using Samtools r982 (Li et al., 2009) ([mpileup-BuDS] filtered by var-filter-snv v1 then false-positive-vcf v1) and annotated by the GMS transcript variant annotator against Ensembl v84. Mutations were filtered to those annotated as nonsynonymous Tier 1 mutations within the protein-coding regions, meeting the following thresholds: minimum 5 reads supporting the variant, minimum 5% variant allele frequency, minimum 20X coverage of the corresponding genomic position in the tumor sample, and minimum 20X coverage of the corresponding genomic position in the normal tail sample (if available). The 8 tail samples were used as a ‘panel of normal’ samples in order to remove common nucleotide polymorphisms and artifacts with the following requirements: SNVs with a minimum 2.5% VAF with 20X coverage detected in at least 2 (25%) of the normal samples or Indels with minimum 2% VAF with 2X coverage detected in at least 2 (25%) of the normal samples. Following this filtering strategy, false positives were manually removed, as previously described (Barnell et al., 2018).

All variants were further filtered by removing those identified in normal mice by the Mouse Genomes Project (v142) (Keane et al., 2011). There were no remaining variants detected in T23, likely indicating that this tumor had low tumor cellularity, making it difficult to differentiate sequencing artifacts from variants detected. There were five tumors (T6, T7, T12, T13, T16, T17) without matched normal tail samples. Recurrent mutations that were identified only in these unmatched tumors were removed as likely germline variants. Furthermore, mutations in olfactory receptor and uncharacterized cDNA regions (*Olfr* and *Rik* genes) were removed.

Copy number variants, including *Kras* amplifications, were identified using CopyCat v0.1 [https://github.com/chrisamiller/copyCat], comparing matched tumor and normal tail samples, if available. In tumors with no matched normal samples, copy number alterations were detected by normalizing the depth of sequencing within each sample over the exons spanning the *Kras* gene locus (in 320-920 bp windows). While there were no variants detected in T23, there was increased normalized coverage over the *Kras* gene locus indicating a possible copy number amplification. Structural variants were predicted using Manta on WGS and WES data (Chen et al., 2016), and intergenic fusions were detected using Integrate (Zhang et al., 2016).

### rPrl expression analysis

A kallisto index was built to incorporate the annotated Ensembl mouse transcriptome (v92) and the full *rPrl* transcript (ENSRNOT00000023412.4) (Zerbino et al., 2018). RNAseq reads were pseudoaligned to this kallisto index to quantify read abundance and expression (TPM) of *rPrl*.

### RNA expression analysis

RNAseq reads were aligned to the mouse reference genome (NCBI build 38, GRCm38) using TopHat. Cufflinks (Trapnell et al., 2010) and HTSeq-count (Anders and Huber, 2010) were used to quantify gene and transcript expression, and differential expression analysis was performed using the DESeq2 R package. The GAGE R package was used for pathway analysis, and data visualization was performed using the ggplot2 R package and GenVisR (Skidmore et al., 2016). Immune infiltrate was assessed by seq-ImmuCC using the Illumina RNAseq platform and the SVR method (Chen et al., 2018). Immune populations were summarized as relative frequencies (the default output of the program).

### RT-PCR

Total RNA was isolated from samples in the Discovery Set (MCPs, AMGs, tumors), WT age-matched (12-14 week) MCPs, and non-tumor mammary cells from aged (15-16 month) mice isolated using the MCP procedure, using the RNeasy Midi Kit (#75142; Qiagen). cDNA was synthesized from 1μg of RNA using M-MLV reverse transcriptase (#M1705; Promega) and random hexamers. Real time PCR was performed using SYBR Green (#4309155; ThermoFisher) on the Applied Biosystems 7300 platform as described (Arendt et al., 2011) using the primer sequences in Table S5. Transcripts of interest were normalized to *18S* RNA, and fold changes calculated using the ΔΔCt method.

## Acknowledgements

K.M.C. was supported by the Cancer Biology Pathway through the Siteman Cancer Center at Washington University School of Medicine. M.G. was supported by the National Human Genome Research Institute (NIH NHGRI R00 HG007940). L.A.S. was supported by the National Institutes of Health (R01 CA157675 and R01 CA179556), and UWCCC Core Grant NIH P30 CA014520. O.L.G. was supported by the National Cancer Institute (NIH NCI K22 CA188163 and U01 CA209936).

## Author Contributions

M.G., L.A.S., K.A.O. and O.L.G. developed the project concept and experimental design. K.M.C. led sequencing experiments and data analysis. K.M.C., E.K.B., Z.L.S., and W.A.M. performed data analysis and prepared figures and tables. K.A.O. and D.E.R. developed methods and carried out mouse experiments and analyses. K.M.C. wrote the manuscript with input from K.K., M.G., K.A.O., L.A.S., and O.L.G.

## Accession Numbers

The reference-aligned whole genome sequence data and sample details for 36 tumor and non-tumor mouse tissues have been submitted to NCBI SRA study XX, BioProject PRJNA489661.

## Supplemental Figures

**Figure S1.**
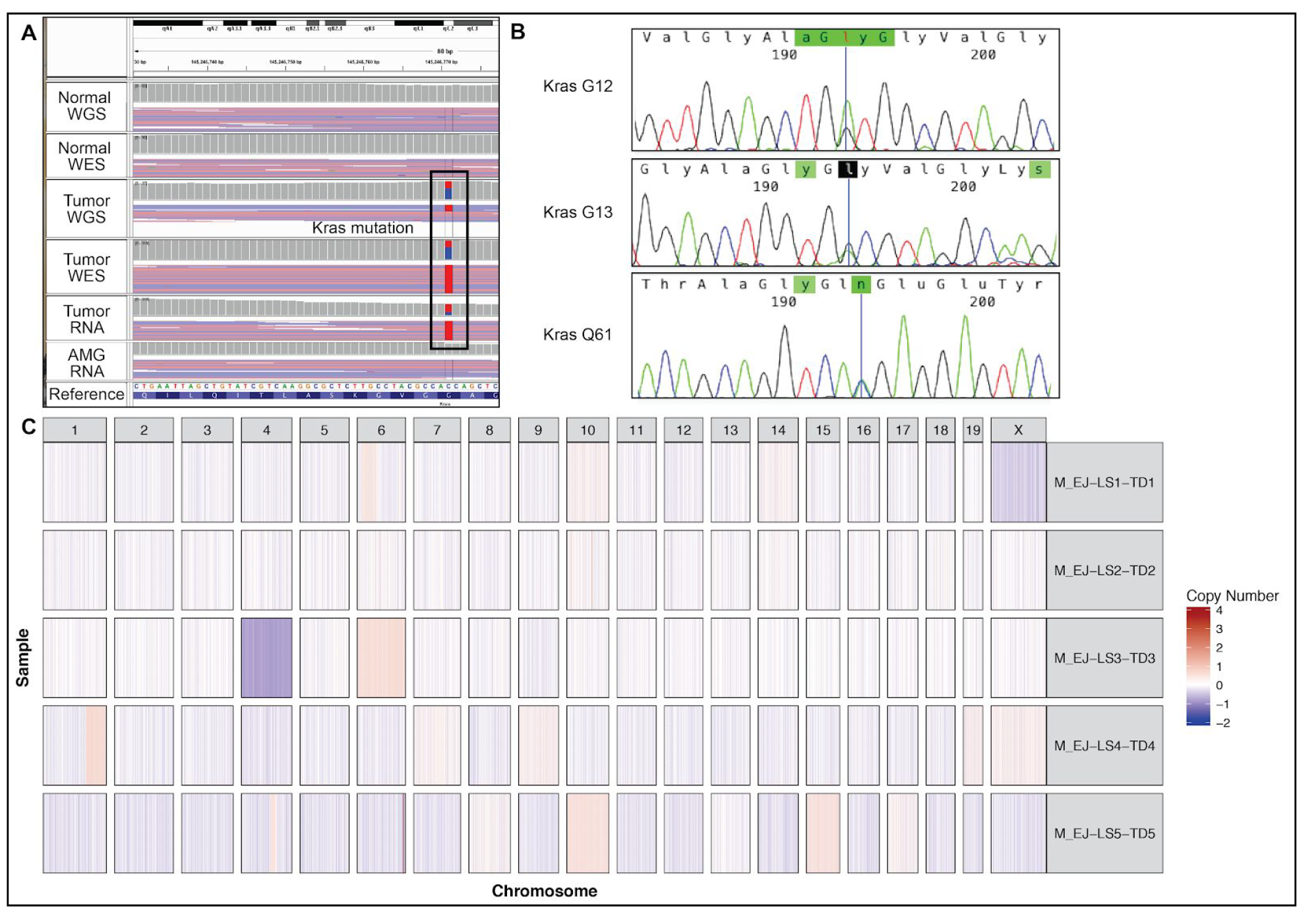
Somatic discovery in the Discovery and Extension Sets. **A.** Representative screenshot from the Integrated Genome Viewer (IGV) displaying a *Kras* G12 mutation in the Discovery Set, present in tumor whole genome, whole exome, and RNA sequencing reads (WGS, WES, RNA), but not in the normal WGS and WES, nor RNA from matched aged mammary glands (AMGs). **B.** Representative Sanger traces from the Extension Set, including mutations detected in *Kras* at either G12, G13, or Q61. **C.** Copy number landscape of the Discovery Set. Fill color represents the relative copy number of the genomic region.

## Supplemental Tables

**Table S1. Sample manifest. Refer to Figure 1.**

This table shows all information corresponding to the samples collected for both Discovery and Extension Sets, including tumors, aged mammary glands, matched normal tails, cell lines, and mammary cell preparations evaluated in this study.

**Table S2. Master variants file of somatic mutations. Refer to Figure 3 and Table 1.**

This table shows all filtered and manually reviewed somatic SNVs and Indels detected in tumors from the Discovery and Extension sets, including variant annotations and read counts corresponding to all WGS, WES, and RNAseq data where available.

**Table S3. Differentially expressed genes. Refer to Figures 4-5.**

DESeq2 was used to evaluate differentially expressed genes across sample types, based upon supervised, pairwise comparisons of gene counts from mammary cell preparations (MCPs), aged mammary glands (AMGs), or tumors (T). Only statistically significant genes (q<0.01) are shown.

**Table S4. Differentially expressed pathways. Refer to Figure 5.**

Upregulated and Downregulated GAGE pathway analysis indicate the supervised, pairwise comparison of mammary cell preparations (MCPs), aged mammary glands (AMGs), or tumors (T) with “same.dir=T” indicating the assumption of directional regulation of genes based upon GO annotations and KEGG pathways. ‘Differentially regulated’ pathways utilized “same.dir=F” for KEGG pathways only. Only statistically significant pathways (q<0.01) are shown.

**Table S5. Primers. Refer to Experimental Procedures.**

Primers used in PCR amplification for Sanger sequencing and RT-PCR experiments.

